# Leptin-receptor neurons in the dorsomedial hypothalamus regulate the timing of circadian rhythms in feeding and metabolism in mice

**DOI:** 10.1101/2020.10.03.324871

**Authors:** Chelsea L. Faber, Jennifer D. Deem, Bao Anh Phan, Tammy P. Doan, Kayoko Ogimoto, Zaman Mirzadeh, Michael W. Schwartz, Gregory J. Morton

## Abstract

Animal behavior and metabolism are tightly coordinated with sleep-wake cycles governed by the brain in harmony with environmental light:dark cycles. Within the brain, the dorsomedial hypothalamic nucleus (DMH) has been implicated in the integrative control of feeding, energy homeostasis, and circadian rhythms [1], but the underlying cell types are unknown. Here, we identify a role for DMH leptin receptor-expressing neurons (DMH^LepR^) in these effects. Using a viral approach, we show that silencing DMH^LepR^ neurons in adult mice not only increases body weight and adiposity, but also shifts circadian rhythms in feeding and metabolism into the light-cycle. Moreover, DMH^LepR^ silencing abolishes the normal increase in dark-cycle locomotor activity characteristic of nocturnal rodents. Furthermore, DMH^LepR^-silenced mice fail to entrain to a restrictive change in food availability. Together, these findings identify DMH^LepR^ neurons as critical determinants of the daily time of feeding and associated metabolic rhythms.

## Introduction

Synchrony between behavior and environmental rhythms enables animals to predict food availability and optimize metabolism in anticipation of daily periods of fasting and feeding [1]. Conversely, mistimed feeding (i.e., food consumption during the normal resting period) impairs metabolism and increases susceptibility to obesity and associated metabolic impairment [2, 3]. While the hypothalamic suprachiasmatic nucleus (SCN) is well-known to entrain circadian rhythmicity in accordance with light:dark cycles, food availability can also entrain metabolic rhythms independently from the SCN [2]. Illustrating this point, although rodents with SCN lesions exhibit profound disruptions in circadian rhythms, they retain the ability to re-train metabolic and behavioral rhythms in accordance with a scheduled meal [3]. Moreover, scheduled feeding has no effect on rhythmic gene expression in the SCN [4], suggesting the existence of extra-SCN food-entrainable oscillators that function to align behavior and metabolism with food availability [1]. Although somewhat controversial [5], evidence suggests the DMH may play such a role [1]. Firstly, the DMH is innervated by the SCN [6], and DMH neurons in turn project to neurons in brain areas regulating metabolism and feeding, including agouti-related protein (AgRP) neurons in the arcuate nucleus (ARC) [7, 8]. Moreover, DMH lesioning in rats not only disrupts circadian rhythms in feeding, locomotion, and core temperature [9, 10], but also precludes entrainment to scheduled feeding [9]. However, the relevant DMH cell types mediating these effects are unknown. Based on recent evidence that DMH neurons expressing leptin receptor (DMH^LepR^) are both sensitive to food availability and make synaptic connections with AgRP neurons to modulate feeding [7], we identified DMH^LepR^ neurons as a candidate population for the circadian control of food intake and associated metabolic rhythms.

## Results and Discussion

### Inactivation of DMH^LepR^ neurons elicits transient hyperphagia and increased adiposity

To determine the role of DMH^LepR^ neurons in feeding and metabolism, we used a viral loss-of-function approach (Figure 1A). Specifically, DMH^LepR^ neurons were permanently silenced following bilateral microinjection of an AAV encoding Cre-dependent tetanus toxin light-chain fused with a GFP reporter (AAV1-CBA-DIO-GFP:TeTx) [11]. Viral transduction was confirmed by histochemical detection of GFP in the DMH (Figure 1B-C); as expected, GFP was undetected in Cre-negative controls (not shown). Outside of the DMH, abundant GFP+ terminals were detected in the ARC (Figure 1B-C), consistent with previous evidence of a DMH^LepR^ → ARC^AgRP^ neurocircuit implicated in feeding control [7, 12].

**Figure 1.**
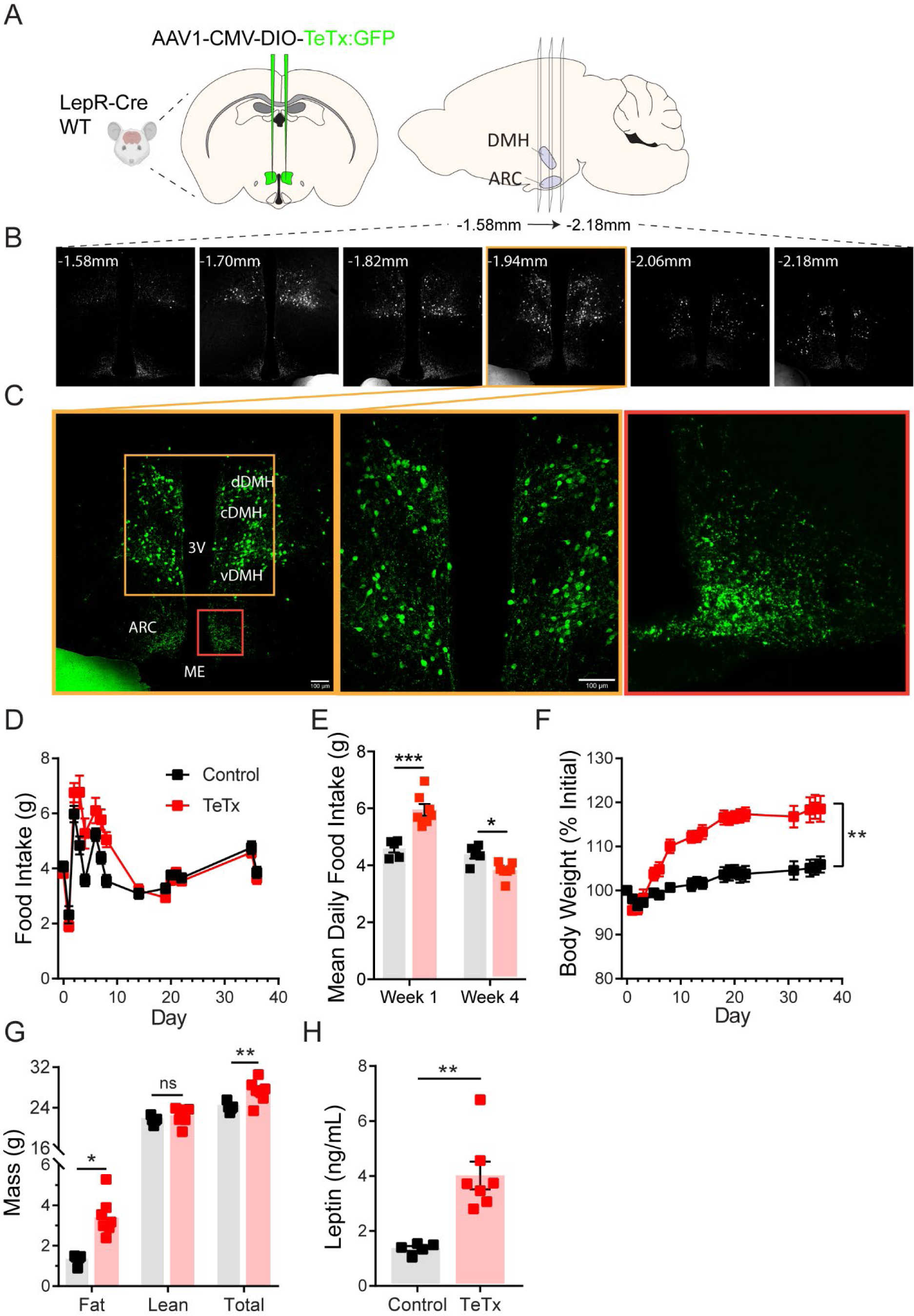
Silencing DMH^LepR^ neurons elicits transient hyperphagia and increased adiposity in adult male mice. (A) Experimental schematic for chronic inhibition of DMH^LepR^ neurons by microinjection on Day 0 of an AAV1 containing a Cre-dependent GFP-fused TeTx delivered bilaterally to the DMH LepR-Cre+ male mice (TeTx; n=7) and Cre-littermate controls (Control; n=7). (B) Stereological fluorescent images from a representative animal showing the rostral-caudal extent of TeTx:GFP expression. (C) *Left*: Colorized, higher magnification view of the boxed orange region from (B). *Middle:* Higher magnification view of the boxed orange region showing neuronal cell bodies targeted within the DMH. *Right*: Higher magnification view of the boxed red region showing TeTx:GFP+ terminals of targeted DMH^LepR^ neurons within the ARC (D) Mean daily food intake following viral microinjection. Two-way ANOVA: F_(1,10)_=4.658; p=0.0563 (main effect of TeTx); F(_14,140)_=4.886; p<0.0001 (time x TeTx interaction). (E) Mean daily food intake from Week 1 relative to Week 4. Two-way ANOVA: F_(1,10)_=5.575; p=0.0399 (main effect of TeTx); F_(1,10)_=39; p<0.001 (time x TeTx interaction). (F) Body weight expressed as %Day 0 value. Two-way ANOVA: F_(1,10)_=20.18; p=0.0012 (main effect of TeTx). F(_19,190)_=14.67; p<0.0001 (time x TeTx interaction). (G) Fat, lean, and total mass 26 days after viral microinjection. Multiple t-tests; t_fat_=4.847; p=0.0014; t_total_=2.884; p=0.016. (H) Plasma leptin 21 days after viral microinjection. Unpaired t-test, t=5.17, p=0.0017. Data are mean ± SEM. For repeated measures, post hoc, Sidak’s test for each time point are indicated on the graph. *p<0.05,**p<0.01, ***p<0.001, ****p<0.0001

Whereas previous evidence showed no effect of acute inhibition of DMH^LepR^ neurons on feeding [7], chronic inactivation of DMH^LepR^ neurons resulted in hyperphagia that was sustained for several days (Figure 1D-E), an effect associated with sustained weight gain (Figure 1F) and a selective increase in adipose mass (Figure 1G), despite daily food intake eventually falling below that of controls (Figure 1E). These effects were accompanied by modestly increased plasma leptin levels (Figure 1H) and elevated fasted levels of both blood glucose (Control vs. TeTx: 72 ± 5.621 vs. 107.1 ± 7.295, t9.969=3.816; p=0.003) and plasma insulin (Control vs. TeTx: 0.49 ± 0.0419 vs. 1.244 ± 0.1229, t6.092=5.807; p=0.001), suggestive of insulin resistance. These findings extend and refine previous work implicating a physiological role for DMH^LepR^ neurons in energy homeostasis [7, 12].

### DMH^LepR^ neurons are required for inhibition of feeding by leptin

As leptin signaling in the DMH has been implicated in the acute anorexic effect of leptin [13], we tested whether DMH^LepR^ inactivation blunts leptin-mediated anorexia. First, the specificity of GFP:TeTx expression in DMH^LepR^ neurons was confirmed by establishing that leptin-induced pSTAT3, a marker for LepR signaling, colocalizes with virally-transduced cells following systemic leptin injection (Figure 2A). Next, control and DMH^LepR^-silenced mice were fasted for 24 h followed by ip injection of either leptin or saline control, after which food was returned. Although control animals lost more weight during the fast (Figure 2B) and exhibited a greater refeeding response following saline-treatment than saline-treated TeTx mice (Figure 2C; dashed bars), the effect of leptin to suppress food intake was absent in DMH^LepR^-silenced mice (Figure 2C; solid bars). These findings extend previous evidence [13] of a key role for DMH^LepR^ neurons in leptin-mediated suppression of fasting-induced refeeding.

**Figure 2.**
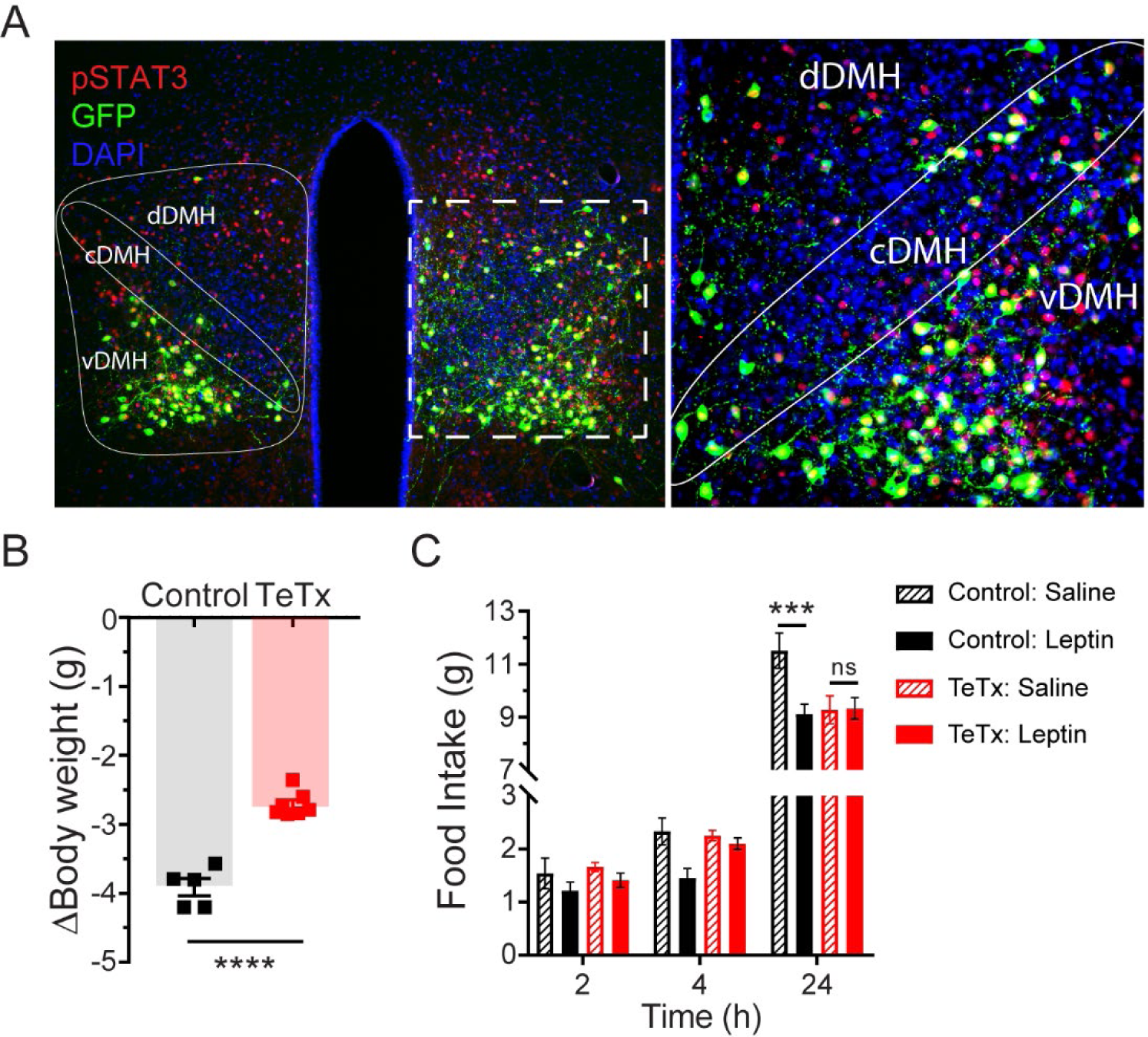
Validation of DMH^LepR^ neuronal targeting and evidence that activation of these neurons is required for leptin-induced anorexia. (A) Left: Representative image showing extensive overlap of pSTAT3 expression in GFP:TeTx-expressing DMH^LepR^ neurons in mice sacrificed 90 minutes following leptin administration (i.p. 3 mg/kg). Right: Higher magnification view of the boxed region from the left. (B) Change in body weight (unpaired T-test, t=8.483, p=0.0001) following a 24h (ZT2 – ZT2’) fast 5 weeks following viral microinjection and before food was returned in (C). (C) Post-fast (24h) refeeding following i.p. injection of saline or leptin (3 mg/kg). Two-way ANOVA: F(1,4)=47.33; p=0.0023 (Controls, main effect of leptin). F(1,6)=0.1203; p=0.7405 (TeTx, main effect of leptin). v-, c-, and dDMH = ventral, central, and dorsal compartments of the dorsomedial hypothalamic nucleus, respectively; 3V = 3rd ventricle; ARC = arcuate nucleus; ME = median eminence. Data are mean ± SEM. For repeated measures, post hoc, Sidak’s test at each time point are indicated on the graph. *p<0.05, ***p<0.001, ****p<0.0001.

### DMH^LepR^ inactivation disrupts diurnal feeding, locomotion, and metabolic rhythms

To determine whether the observed impairments in energy homeostasis were associated with changes in circadian rhythmicity, we obtained continuous measures of energy intake, energy expenditure, and locomotor activity using indirect calorimetry. We found that unlike control mice, which exhibited typical nocturnal feeding behavior, the phase of food intake was shifted in DMH^LepR^-silenced mice (Figure 3A), such that dark-cycle food intake was decreased and light-cycle intake increased (Figure 3B). Similarly, while control mice displayed a typical increase in dark-cycle locomotor activity, this was absent in DMH^LepR^-silenced mice (Figure 3C-D). Rhythms in other metabolic parameters were similarly shifted and blunted by DMH^LepR^ inactivation. Specifically, we found that heat production in DMH^LepR^-silenced mice was reduced selectively in the dark cycle (Figure 3E-F) and respiratory-exchange ratio (RER) was elevated in the light cycle (Figure 3G-H), indicative of an increase in carbohydrate utilization consistent with the increased feeding during this time (Figure 3A-B). Together, these findings identify DMH^LepR^ neuron activity as a crucial determinant of appropriately timed circadian rhythms in feeding, locomotor activity, and associated metabolic parameters.

**Figure 3.**
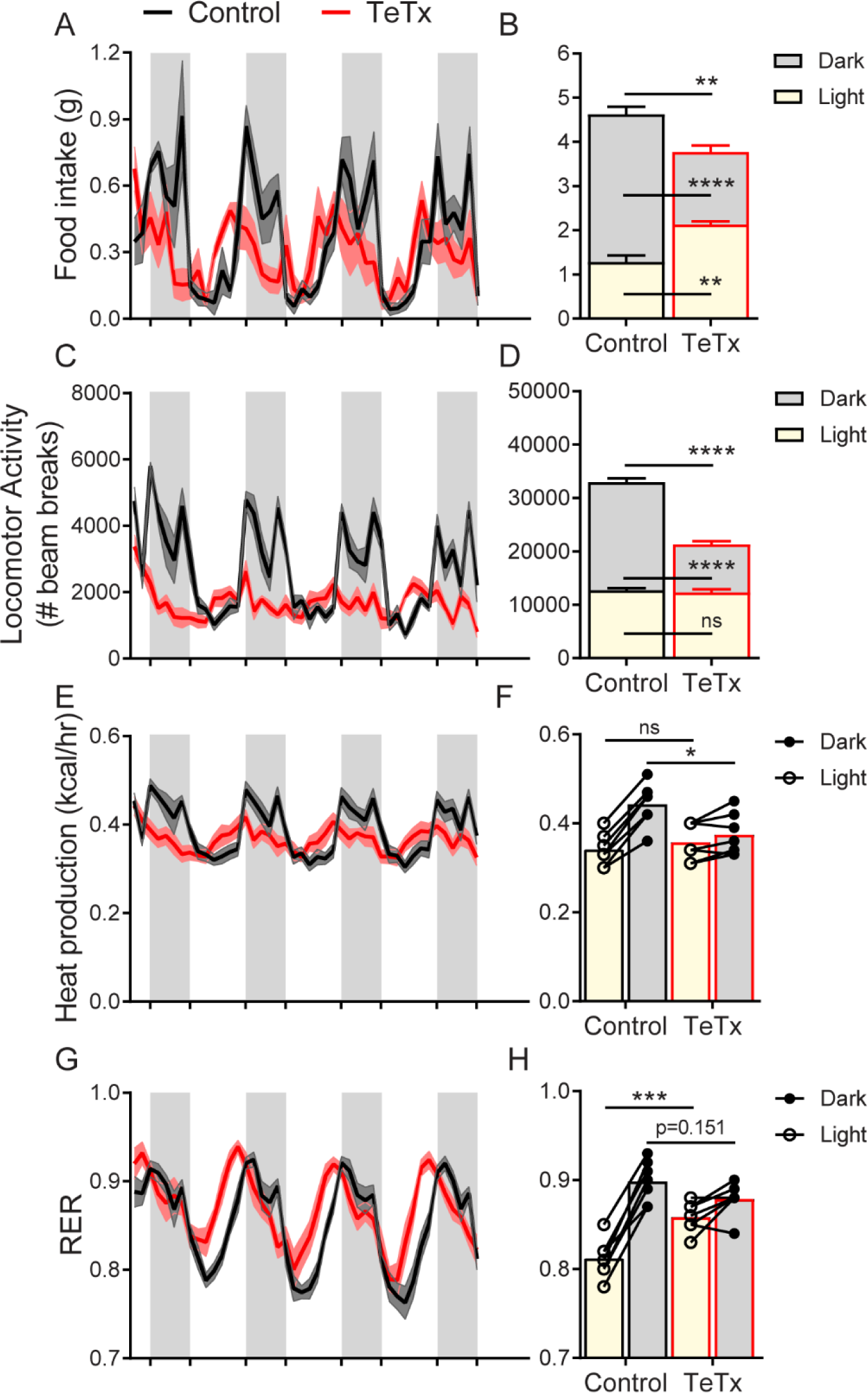
DMH^LepR^ neuron inactivation disrupts circadian patterns of food intake, locomotor activity, heat production, and substrate utilization. 2h-binned continuous measures (left panels) and mean values across the light (L) and dark (D) periods (right panels) 30 days following microinjection of TeTx:GFP (TeTx; n=7) or GFP control (Control; n=7) to the DMH of LepR-Cre+ male mice. (A) Food intake. Two-way ANOVA: F(1,12)=12; p=0.0047 (main effect of TeTx). F(87,1044)=2.354; p<0.0001 (time x TeTx interaction). (B) Mean food intake from A during L, D, and 24h-period. Two-way ANOVA: F(1,12)=9.567; p=0.0093 (main effect of TeTx). (C) Locomotor activity. Two-way ANOVA: F(1,12)=93.22; p<0.0001 (main effect of TeTx). (D) Mean locomotor activity from C during L, D, and 24h-period. Two-way ANOVA: F(1,12)=110.4; p<0.0001 (main effect of TeTx). (E) Heat production. Two-way ANOVA: F(1,12)=1.006; p=0.3357 (main effect of TeTx). (F) Mean heat production from E during L and D periods. Two-way ANOVA: F(1,12)=1.209; p=0.2930 (main effect of TeTx). (G) Respiratory exchange ratio (RER). Two-way ANOVA: F(1,12)=2.789; p=0.1208 (main effect of TeTx). (H) Mean RER from G during L and D periods. Two-way ANOVA: F(1,12)=2.04; p=0.1788 (main effect of TeTx). Data are mean ± SEM. For repeated measures, post hoc, Sidak’s test at each time point are indicated on the graph. *p<0.05,**p<0.01, ***p<0.001, ****p<0.0001.

### Female DMH^LepR^-silenced mice recapitulate weight gain and circadian disruption seen in males

We also tested whether the phenotype is conserved between sexes. Although female DMH^LepR^-silenced mice did not exhibit the transient hyperphagia observed in males (Supplemental Figure 2A-B), they nonetheless developed mild obesity (Supplemental Figure 2C-D). Females also exhibited disrupted circadian rhythms in food intake (Supplemental Figure 3A-B), locomotor activity (Supplemental Figure 3C-D), heat production (Supplemental Figure 3E-F), and RER (Supplemental Figure 3G-H) similar to those observed in male DMH^LepR^-silenced mice. The key role for DMH^LepR^ neurons in circadian behavioral and metabolic control identified in males, therefore, extends to females as well. Given that, compared to male mice [14], female mice are protected from both hyperphagia and disrupted circadian rhythms with HFD [15], future studies are warranted to determine whether sensitivity to HFD is intact in both male and female mice with DMH^LepR^ inactivation and if the DMH lies downstream of circuits mediating sexually-dimorphic responses to HFD.

### Silencing DMH^LepR^ neurons prevents behavioral adaptation to restricted feeding

To determine the extent to which circadian disruptions in metabolism in DMH^LepR^-silenced mice are secondary to the shift in daily patterns of food intake and whether DMH^LepR^ neurons are required to entrain feeding behavior, a time-restricted feeding (TRF) paradigm was implemented.

We restricted food availability to the dark-cycle, active period (ZT14-ZT24) in both DMH^LepR^-silenced and control mice. After a 5-day TRF acclimation period, both groups were subjected to indirect calorimetry for 5 days during TRF followed by 3 days of *ad libitum* feeding (Figure 4A).

**Figure 4.**
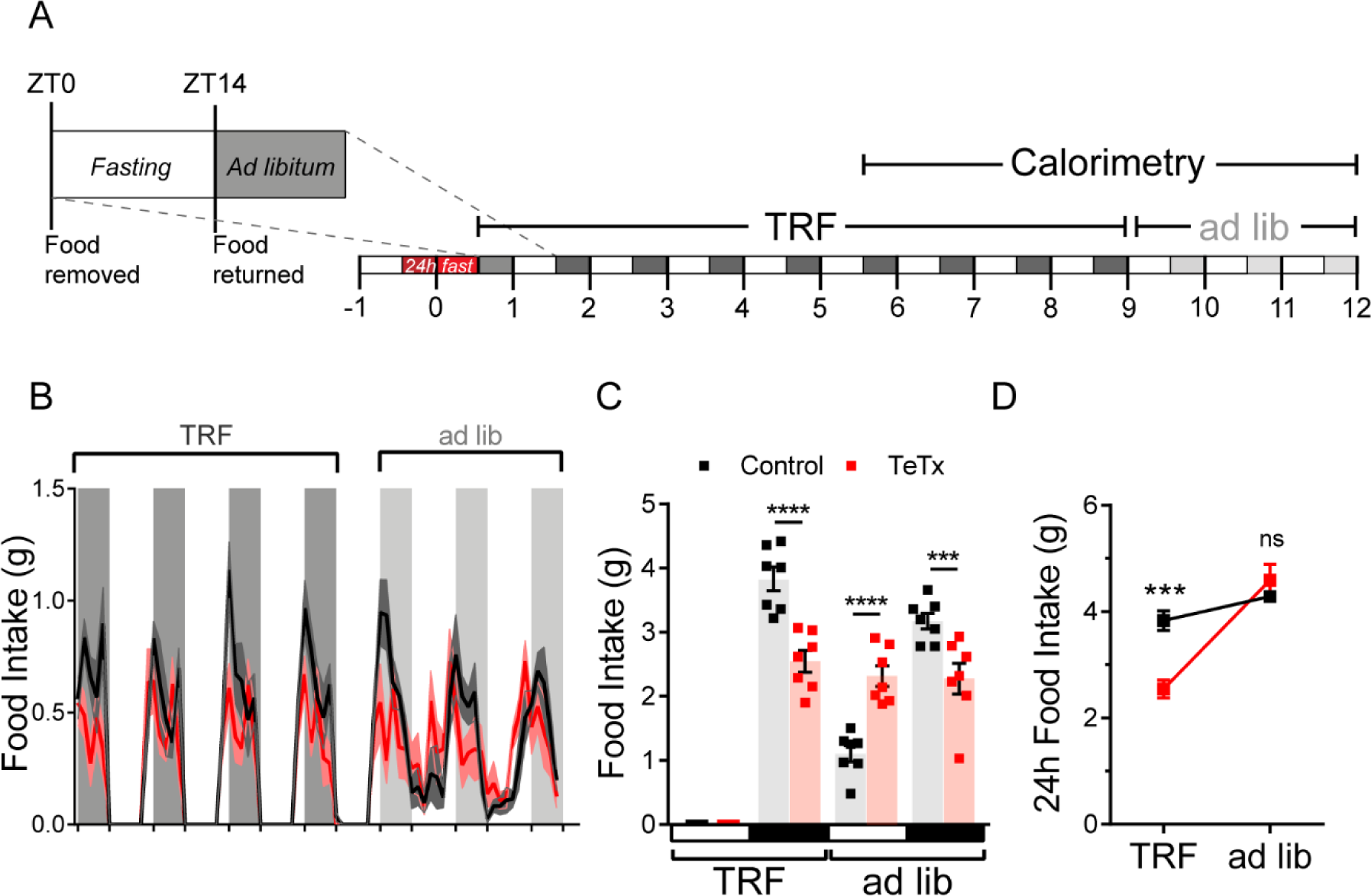
DMH^LepR^ neurons are required for adaptation to a dark-cycle restricted feeding schedule. (A) Experimental timeline. 6 weeks following microinjection of GFP:TeTx (TeTx; n=7) or GFP control (Control; n=7) to the DMH of LepR-Cre+ male mice, mice were acclimated to time-restricted feeding (TRF) in their home cages for a 5 day lead-in before transfer into direct calorimetry. TRF was maintained in calorimetry for an additional 4 days, followed by ad libitum (ad lib) feeding. (B) 2h-binned continuous measures of food intake during TRF and transition back to ad lib feeding. (C) Mean L:D food intake from B under TRF and ad lib feeding. Two-way ANOVA: F_(1,12)_=5.084; p=0.0436 (main effect of TeTx); F_(3,36)_=27.91; p<0.0001 (time x TeTx interaction). (D) Mean 24h-period food intake from C during TRF and ad lib feeding. Two-way ANOVA: F_(1,12)_=5.097; p=0.0434 (main effect of TeTx); F_(1,12)_=47.8; p<0.0001 (main effect of TRF); F_(1,12)_=19.58; p=0.0008 (TRF x TeTx interaction). Within treatment comparison (TRF vs ad lib): Control t_(12)_=1.759; p=0.1971; TeTx t_(12)_=8.018; p<0.0001. Data are mean ± SEM. For repeated measures, post hoc, Sidak’s test at each time point are indicated on the graph. *p<0.05,**p<0.01, ***p<0.001, ****p<0.0001.

During TRF acclimation, body weight oscillated daily as expected in both groups, being higher after food was available during the dark cycle, and lower after light-cycle fasting. However, unlike control mice which were able to maintain their weight during TRF, DMH^LepR^-silenced mice exhibited a small reduction in body weight (Supplemental Figure 4B), likely because control mice compensated for the imposed light-cycle fast by increasing dark-cycle food intake unlike the DMH^LepR^-silenced mice (Figure 4B-D). Upon restoration of *ad libitum* feeding, DMH^LepR^-silenced mice exhibited rebound hyperphagia sufficient to recover lost weight (Figure 4B-D). Interestingly, this hyperphagic response was limited to the light cycle, as DMH^LepR^-silenced mice rapidly reverted to their mistimed feeding rhythms (Figure 4B-C). Together, these findings indicate that DMH^LepR^ neuron activity is required to entrain feeding behavior during dark-cycle TRF. Although mechanisms underlying this adaptive response await further study, the capacity to increase intake when food is available for a restricted window each day requires the ability to anticipate when food will be available in association with a variety of metabolic and neuroendocrine adaptations, e.g., [16]. Our findings also reveal that although DMH^LepR^-silenced mice are capable of mounting rebound hyperphagia following weight loss, this response appears to require *ad libitum* access to food during the light-cycle, a time when normal mice consume little food.

### Conclusion

Our work identifies a crucial physiological role for DMH^LepR^ neurons in circadian regulation of feeding behavior, locomotion, and associated metabolic parameters. Activity of these neurons is also necessary to adapt feeding during a restricted feeding paradigm. Given evidence from both humans and rodents that mistimed feeding can predispose to obesity and T2D [17, 18], these findings have relevance to the pathogenesis of both disorders. An improved understanding of the neural circuits underlying endogenous rhythms of behavior, feeding, and metabolism may facilitate the development of new therapeutic and dietary strategies for the treatment of obesity and related metabolic disorders in humans.

## Research design and methods

### Mice

All procedures were performed in accordance with the National Institutes of Health Guide for the Care and Use of Laboratory Animals and were approved by the Animal Care Committee at the University of Washington. Following stereotaxic surgery, all studied animals were individually housed with ad libitum access to standard chow diet (LabDiet 5053) in a temperature and humidity-controlled facility with 14:10 light:dark cycles. Adult *Lepr*^*IRES-Cre/+*^ (LepR-Cre) mice (Jackson Laboratory no. 008320) were used for all experiments, unless otherwise noted.

### Stereotactic Surgeries

The viral vector AAV1-CBA-DIO-GFP:TeTx (TeTx) was generated as described [19], and generously provided by Dr. Richard Palmiter and Dr. Larry Zweifel (University of Washington, Seattle, WA). For viral microinjection, animals were placed in a stereotaxic frame (Kopf 1900; Cartesian Research Inc., Tujunga, CA) under isoflurane anesthesia. The skull was exposed with a small incision, and two small holes were drilled for bilateral 200-nL injection volume of TeTx into the DMH of LepR-Cre or Cre-negative littermate mice based on coordinates from the Mouse Brain Atlas [20]: anterior-posterior (AP) −1.6, dorsal-ventral (DV) −5.6 mm, and lateral 0.40 mm. Adeno-associated virus (AAV) was delivered using a Hamilton syringe with a 33-gauge needle at a rate of 50 nL/min (Micro4 controller), followed by a 5-min wait at the injection site and a 1-min wait 0.05 mm dorsal to the injection site before needle withdrawal. Animals received a perioperative subcutaneous injection of buprenorphine hydrochloride (0.05 mg/kg) (Reckitt Benckiser, Richmond, VA). Viral expression was verified post hoc in all animals, and any data from animals in which the virus expressed outside the targeted area were excluded from the analysis.

### Body Composition Analysis

Measurements of body lean and fat mass were determined in live, conscious mice by use of quantitative magnetic resonance spectroscopy (QMR; EchoMRI-700TM; Echo MRI, Houston, TX) by the University of Washington Nutrition Obesity Research Center (NORC) Energy Balance Core.

### Leptin effects on food intake and pSTAT3-induction

To validate the ability of leptin to elicit pSTAT3 signaling in DMH^LepR^ neurons, ad lib fed mice were injected intraperitoneally with leptin (5 mg/kg; Dr. Parlow; National Hormone Peptide Program) and perfused 90 min later, as described below.

To assess the ability of leptin to suppress the compensatory hyperphagia that normally follows a prolonged fast, mice were fasted for 24 h from ZT2 – ZT2’. On the second day, leptin (3 mg/kg) or vehicle-control (PBS, pH 7.9) was injected intraperitoneally in mice 15 min before preweighed food was placed back in the cage, and intake was monitored for the following 24 h.

### Indirect Calorimetry, Food Intake, and Activity

Mice were acclimated to calorimetry cages prior to study and data collection. Energy expenditure measurements were obtained by a computer-controlled indirect calorimeter System (Promethion, Sable Systems, Las Vegas NV) with support from the Energy Balance Core of the NORC at the University of Washington, as previously described [21]. Oxygen consumption (VO2) and carbon dioxide production (VCO2) were measured for each mouse for 1-min at 10-min intervals, and food and water intakes were measured continuously while mice were housed in a temperature- and humidity-controlled cabinet (Caron Products and Services, Marietta, OH NV). Ambulatory activity was determined simultaneously and beam breaks in the x-, y- and z-axes were scored as an activity count, and a tally was recorded every 10 min. Data acquisition and instrument control were coordinated by MetaScreen v.1.6.2, and raw data were processed using ExpeData v.1.4.3 (Sable Systems, Las Vegas, NV) using an analysis script documenting all aspects of data transformation.

### Time Restricted Feeding (TRF)

Food was removed each morning at the start of the light cycle (ZT0) and returned at the start of the dark cycle (ZT14); body weight was also measured at both ZT0 and ZT14 daily. To eliminate the initial effects of varying fed status of animals, 1 day before TRF animals were fasted for 24 h from ZT14 (on Day −1) to ZT14’ (on Day 0) before TRF began. Animals were then subjected to indirect calorimetry for 5 additional days during TRF before returning to ad lib feeding for the remaining 3 days of study (Figure 4A).

### Immunohistochemistry

For brain immunohistochemical (IHC) analyses, animals were terminally anesthetized with ketamine:xylazine and transcardially perfused with phosphate-buffered saline (PBS) followed by 4% paraformaldehyde (PFA) in 0.1 mol/L PBS. Brains were removed and postfixed overnight, then transferred into 30% sucrose overnight or until brains sunk in solution. Brains were subsequently sectioned on a freezing-stage microtome (Leica) to obtain 30*μ*m coronal sections in four series. A single series of sections per animal was used in histological studies, and the remainder stored in −20 °C in cryoprotectant. Brain sections were washed in PBS with Tween-20, pH 7.4 (PBST) overnight at 4C. Sections were then washed at room temperature in PBST (3×8 min), followed by a blocking buffer (5% normal donkey serum (NDS), 1% bovine serum albumin (BSA) in PBST with azide) for 60 minutes with rocking. Sections were then incubated overnight at 4C in blocking buffer containing primary antiserum (goat anti-GFP, Fitzgerald, 1:1000; rabbit anti-pSTAT3, Sigma-Aldrich, St Louis, Missouri, 1:1000). Next, sections were washed (3 x 8 min) in PBST before incubating in secondary donkey anti-goat IgG Alexa 488 (Jackson ImmunoResearch Laboratories, West Grove, PA) diluted 1:1000 in blocking buffer. Sections were washed (3 x 8 min) in PBST before incubating with DAPI for 8 minutes, followed by a final wash (3 x 10 min) in PBS. Sections were mounted to slides and imaged using a Leica SP8X confocal.

### Tissue Processing, Blood Collection

Tail blood for plasma hormonal measurement was collected at indicated times. Blood was collected via EDTA-coated capillary tubes and centrifuged at 4 °C (7,000 rpm, 4 min) and plasma was subsequently removed and stored at −80 °C for subsequent assay. Plasma leptin (Crystal Chem, Elk Grove Village, IL; #90030) and plasma insulin (Crystal Chem, Elk Grove Village, IL; #90080) were determined by ELISA.

### Statistical Analyses

All results are presented as means ± SEM. *P* values for unpaired comparisons were calculated by two-tailed Student’s *t* test. Time course comparisons between groups were analyzed using a two-way repeated measures ANOVA with main effects of treatment (control vs. TeTx) and time. All post hoc comparisons were determined using Sidak’s correction for multiple comparisons. All statistical tests indicated were performed using Prism (version 7.4; GraphPad, CA) software.

## Acknowledgements

We thank R. Palmiter, C. Campos, L. Zweifel, and M. Baird for producing the TeTx virus, J. T. Nelson for assistance with metabolic experiments, and V. Damian for maintaining the mouse colony. We also thank R. Palmiter for editing the manuscript. We are grateful to N. Peters at the University of Washington Keck Imaging Center for technical assistance and the National Institutes of Health (S10-OD-016240) for support to the W.M. Keck Foundation Center for Advanced Studies in Neural Signaling. This work was supported by NIH grants F31-DK113673 (C.L.F.), T32-GM095421 (C.L.F.), DK089056 and DK124238 (G.J.M.), DK083042 and DK101997 (M.W.S.); the NIDDK-funded Nutrition Obesity Research Center (DK035816) and Diabetes Research Center (DK017047) and the Diabetes, Obesity and Metabolism (T32 DK007247; C.L.F.) and Nutrition, Obesity and Atherosclerosis (T32 HL007028; J.D.D.) Training Grants at the University of Washington; a Dick and Julia McAbee Endowed Fellowship (J.D.D.); an American Diabetes Association Innovative Basic Science Award (ADA 1-19-IBS-192) (G.J.M.); and an American Diabetes Association Fellowship Grant (ADA 1-19-PDF-103) (J.D.D.).

## Author Contributions

C.L.F. conceived and designed research studies, performed stereotaxic surgeries, acquired and analyzed data, and wrote and edited the manuscript. J.D.D., B.A.P., and T.P.D. provided technical assistance with histology and longitudinal animal monitoring. K.O. performed calorimetry experiments. Z.M., M.W.S., and G.J.M. provided guidance and resources and revised the manuscript. All authors approved the final version of the manuscript.

