## Supplemental Figures for "Leptin-receptor neurons in the dorsomedial hypothalamus regulate the timing of circadian rhythms in feeding and metabolism in mice"

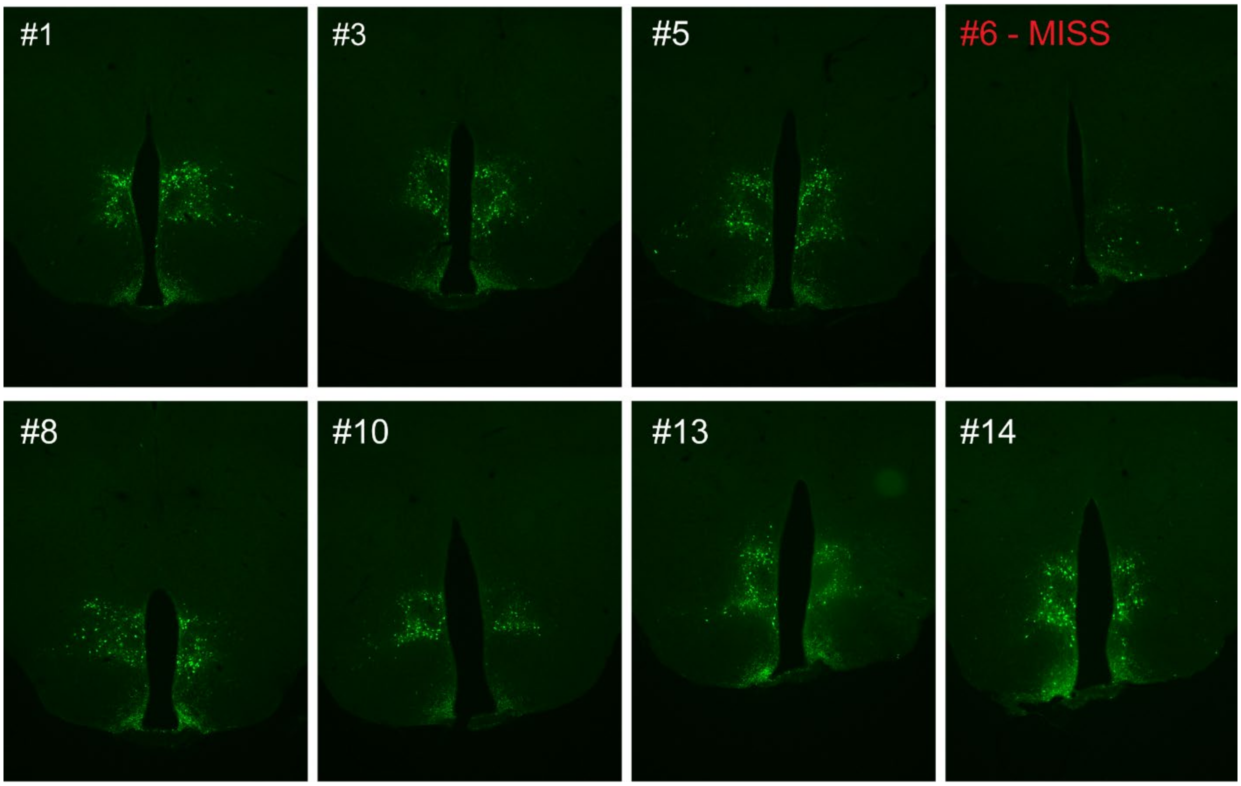


**Supplemental Figure 1. Representative viral expression is evident in both the ventral and dorsal compartments of the DMH following microinjection of TeTx to the DMH of LepR-Cre+ male mice.**
Animal #6 represents a surgical miss and was excluded from all analyses.

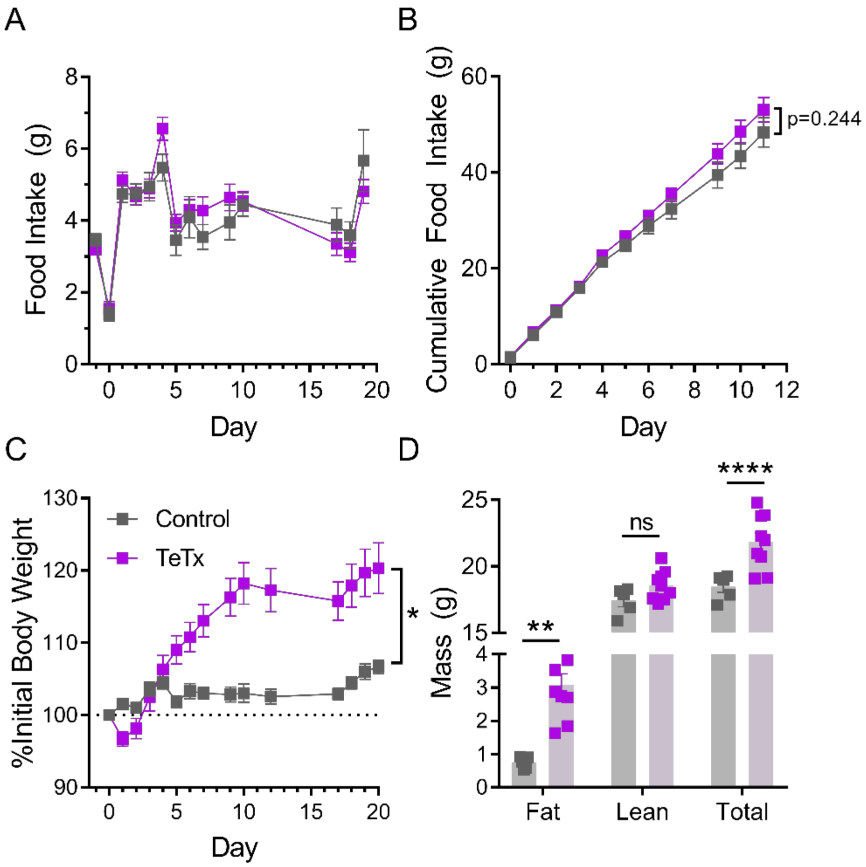


**Supplemental Figure 2 Inhibition of DMH$^{LepR}$ neurons in female mice recapitulates body weight and fat mass increase observed in males, but not acute hyperphagia.**

(A) Average daily food intake following chronic inhibition of DMH$^{LepR}$ neurons by bilateral microinjection on Day 0 of a cre-dependent TeTx:GFP delivered to LepR-Cre+ female mice (TeTx; n=9) and Cre- littermate controls (Control; n=5). Two-way ANOVA: $F_{(1,12)}$=0.3474; p=0.5665 (main effect of TeTx); $F_{(13,156)}$=1.563, p=0.1014 (time x TeTx interaction).

(B) Cumulative food intake for the first 11 days in A (inset). Two-way ANOVA: $F_{(1,12)}$=1.503; p=0.2437 (main effect of TeTx); $F_{(10,120)}$=1.427, p=0.1766 (time x TeTx interaction).

(C) Daily body weight expressed as %Day 0 value. Two-way ANOVA: $F_{(1,12)}$=7.11; p=0.0205 (main effect of TeTx); $F_{(14,168)}$=15.83, p<0.0001 (time x TeTx interaction).

(D) Fat, lean, and total mass 14 days after viral microinjection. Multiple t-tests; $t_{fat}$=3.268; p=0.0024; $t_{total}$=4.705; p<0.0001.

Data are mean ± SEM. *p<0.05,**p<0.01, ***p<0.001, ****p<0.0001.

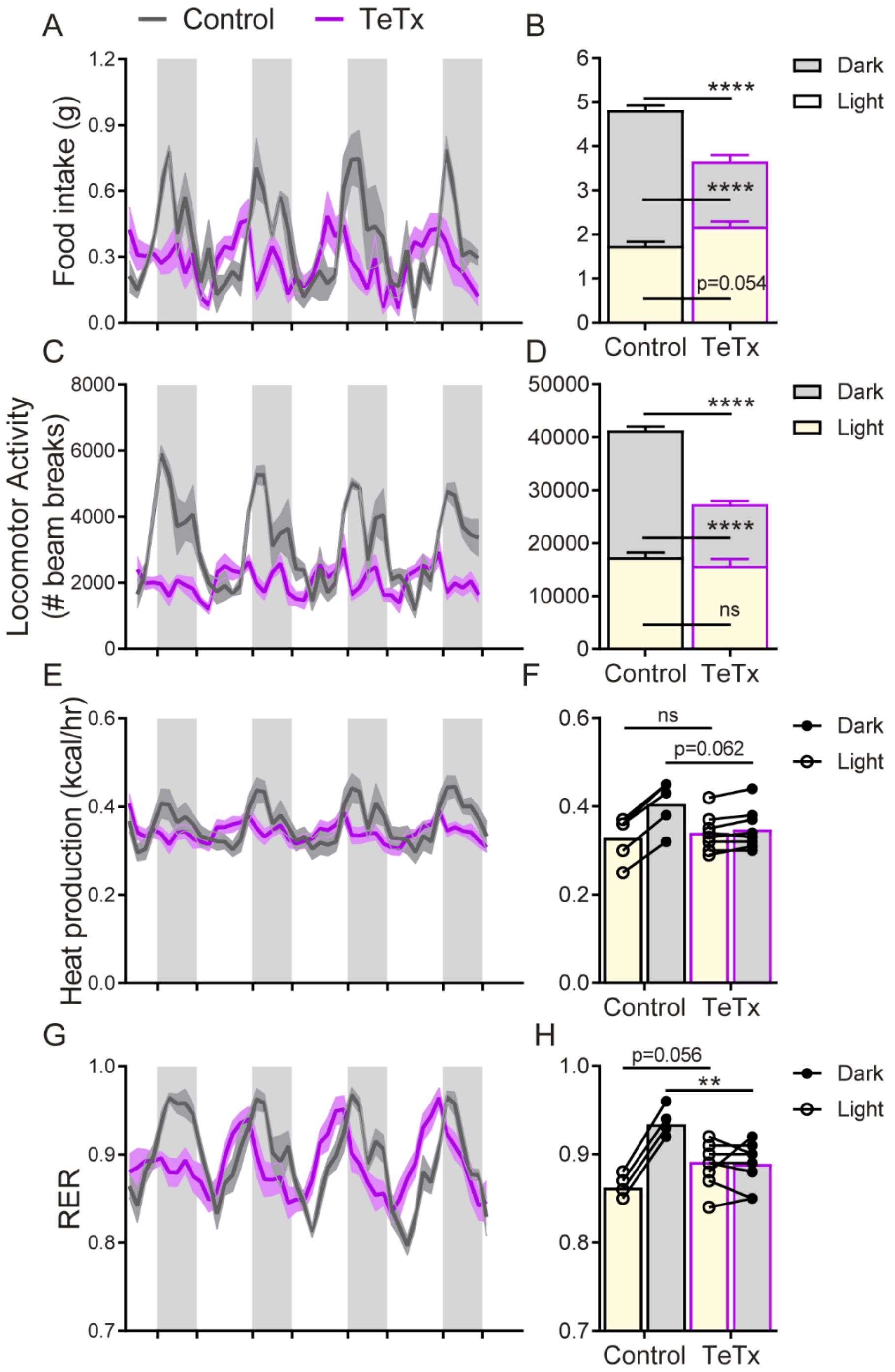


**Supplemental Figure 3 Inhibition of DMH$^{LepR}$ neurons in female mice recapitulates the effect in males to disrupt circadian rhythms.**

2h-binned continuous measures (left panels) and mean values across the light (L) and dark (D) periods (right panels) 30 days following microinjection of TeTx:GFP (TeTx; n=9) or GFP control (Control; n=5) to the DMH of LepR-Cre+ female mice.

(A) Food intake. Two-way ANOVA: $F_{(1,12)}=12.32$; p=0.0043 (main effect of TeTx). $F_{(89,1068)}=2.766$, p<0.0001; $F_{(89,1068)}=2.766$; p<0.0001 (time x TeTx interaction).

(B) Mean food intake from A during L, D, and 24h-period. Two-way ANOVA: $F_{(1,12)}=16.26$; p=0.0017 (main effect of TeTx).

(C) Locomotor activity. Two-way ANOVA: $F_{(1,12)}$=36.22; p<0.0001 (main effect of TeTx); $F_{(89,1068)}$=5.197; p<0.0001 (time x TeTx interaction).
(D) Mean locomotor activity from C during L, D, and 24h-period. Two-way ANOVA: $F_{(1,12)}$=27.98; p=0.0002 (main effect of TeTx).
(E) Heat production. Two-way ANOVA: $F_{(1,12)}$=0.5405; p=0.4764 (main effect of TeTx); $F_{(89,1068)}$=5.903; p<0.0001 (time x TeTx interaction).

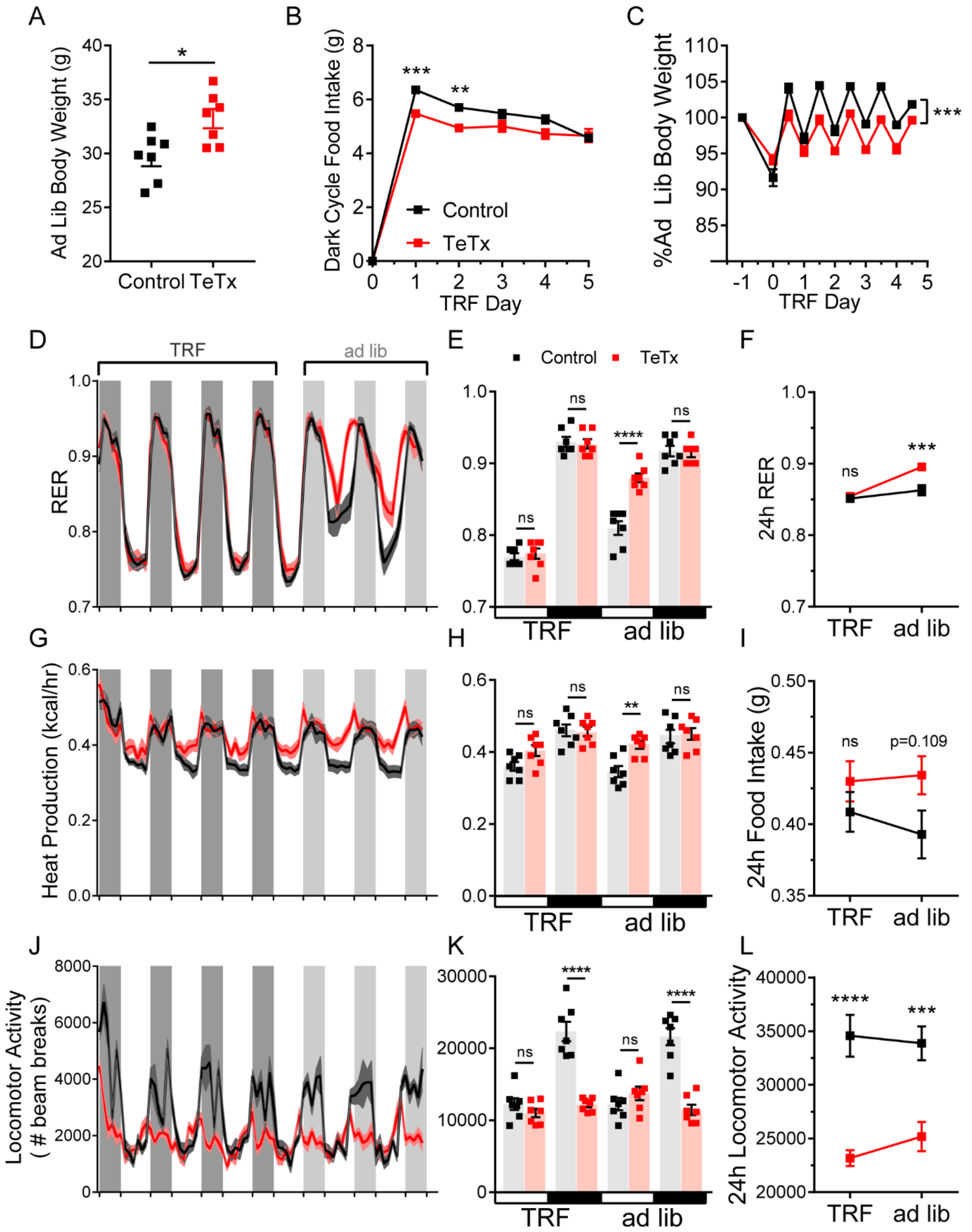


**Supplemental Figure 4 TRF corrects phase-shifts in RER, but has no effect on locomotor activity (associated with Figure 4).**

(A) Ad lib body weight before TRF paradigm. Unpaired t-test, $t_{(11.92)}$=2.946, p=0.0123.

(B) Dark-cycle food intake during TRF lead-in. Two-way ANOVA: $F_{(1,12)}$=10.74; p=0.0066 (main effect of TeTx).

(C) Body weight during TRF lead-in expressed as a percentage of ad lib (pre-TRF) body weight in B. Two-way ANOVA: $F_{(1,12)}$=19.49; p=0.0008 (main effect of TeTx).
(D) 2h-binned continuous measures of respiratory exchange ratio (RER) during TRF and transition back to ad lib feeding.
(E) Mean L:D RER from D during TRF and ad lib feeding. Two-way ANOVA: $F_{(1,12)}$=6.878; p=0.0223 (main effect of TeTx); $F_{(3,36)}$=17.84; p<0.0001 (time x TeTx interaction).
(F) Mean 24h-period RER from E during TRF and ad lib feeding. Two-way ANOVA: $F_{(1,12)}$=9.973; p=0.0083 (main effect of TeTx); $F_{(1,12)}$=28.13; p=0.0002 (main effect of TRF); $F_{(1,12)}$=9.062; p=0.0109 (TRF x TeTx interaction). Within treatment comparison (TRF vs ad lib): Control $t_{(12)}$=1.622; p=0.2445; TeTx $t_{(12)}$=5.879; p=0.0001.
(G) 2h-binned continuous measures of heat production during TRF and transition back to ad lib feeding.
(H) Mean L:D heat production from G during TRF and ad lib feeding. Two-way ANOVA: $F_{(1,12)}$=2.486; p=0.1408 (main effect of TeTx); $F_{(3,36)}$=14.8; p<0.0001 (time x TeTx interaction).
(I) Mean 24h-period heat production from H during TRF and ad lib feeding. Two-way ANOVA: $F_{(1,12)}$=2.559; p=0.1357 (main effect of TeTx); $F_{(1,12)}$=0.8136; p=0.3848 (main effect of TRF); $F_{(1,12)}$=2.492; p=0.1404 (TRF x TeTx interaction). Within treatment comparison (TRF vs ad lib): Control $t_{(12)}$=1.754; p=0.1988; TeTx $t_{(12)}$=0.4783; p=0.8711.
(J) 2h-binned continuous measures of locomotor activity during TRF and transition back to ad lib feeding.
(K) Mean L:D locomotor activity from J during TRF and ad lib feeding. Two-way ANOVA: $F_{(1,12)}$=27.83; p=0.0.0002 (main effect of TeTx); $F_{(3,36)}$=38.22; p<0.0001 (time x TeTx interaction).
(L) Mean 24h-period locomotor activity from K during TRF and ad lib feeding. Two-way ANOVA: $F_{(1,12)}$=27.83; p=0.0002 (main effect of TeTx); $F_{(1,12)}$=0.5965; p=0.4549 (main effect of TRF); $F_{(1,12)}$=2.562; p=0.1354 (TRF x TeTx interaction). Within treatment comparison (TRF vs ad lib): Control $t_{(12)}$=0.5857; p=0.8142; TeTx $t_{(12)}$=1.678; p=0.2242.
Data are mean ± SEM. For repeated measures, post hoc, Sidak's test at each time point are indicated on the graph. *p<0.05,**p<0.01, ***p<0.001, ****p<0.0001.

15. Palmisano BT, Stafford JM, Pendergast JS (2017) High-Fat feeding does not disrupt daily rhythms in female mice because of protection by ovarian hormones. Front Endocrinol (Lausanne) 8(MAR):1–11. https://doi.org/10.3389/fendo.2017.00044
16. Drazen DL, Vahl TP, D'Alessio DA, Seeley RJ, Woods SC (2006) Effects of a Fixed Meal Pattern on Ghrelin Secretion: Evidence for a Learned Response Independent of Nutrient Status. Endocrinology 147(1):23–30. https://doi.org/10.1210/en.2005-0973
17. Challet E (2019) The circadian regulation of food intake. Nat Rev Endocrinol 15(7):393–405. https://doi.org/10.1038/s41574-019-0210-x
18. Huang W, Ramsey KM, Marcheva B, Bass J (2011) Circadian rhythms, sleep, and metabolism. J Clin Invest 121(6):2133–2141. https://doi.org/10.1172/JCI46043.rons
19. Campos CA, Bowen AJ, Schwartz MW, Palmiter RD (2016) Parabrachial CGRP Neurons Control Meal Termination. Cell Metab 23(5):811–820. https://doi.org/10.1016/j.cmet.2016.04.006
20. Franklin KBJ, Paxinos G (2008) The mouse brain in stereotaxic coordinates. Acad Press
21. Kaiyala KJ, Ogimoto K, Nelson JT, Schwartz MW, Morton GJ (2015) Leptin signaling is required for adaptive changes in food intake, but not energy expenditure, in response to different thermal conditions. PLoS One 10(3):1–19. https://doi.org/10.1371/journal.pone.0119391
